# An in silico FSHD muscle fibre for modelling DUX4 dynamics and predicting the impact of therapy

**DOI:** 10.1101/2022.12.12.520053

**Authors:** Matthew V. Cowley, Johanna Pruller, Massimo Ganassi, Peter S. Zammit, Christopher R. S. Banerji

**Author notes:** M. V. Cowley and J. Pruller should be considered joint first authors. Corresponding Author: C.R.S Banerji.

## Abstract

Facioscapulohumeral muscular dystrophy (FSHD) is an incurable myopathy linked to over-expression of the myotoxic transcription factor DUX4. Targeting DUX4 is the leading therapeutic approach, however it is only detectable in 0.1-3.8% of FSHD myonuclei. How rare DUX4 drives FSHD and the optimal anti-DUX4 strategy is unclear. We combine stochastic gene expression with compartment models of cell states, building a simulation of DUX4 expression and consequences in FSHD muscle fibres. Investigating iDUX4 myoblasts, scRNAseq and snRNAseq of FSHD muscle we estimate parameters including DUX4 mRNA degradation, transcription and translation rates and DUX4 target gene activation rates. Our model accurately recreates the distribution of DUX4 and target gene positive cells seen in scRNAseq of FSHD myocytes. Importantly we show DUX4 drives significant cell death despite expression in only 0.8% of live cells. Comparing scRNAseq of unfused FSHD myocytes to snRNAseq of fused FSHD myonuclei, we find evidence of DUX4 protein syncytial diffusion and estimate its rate via genetic algorithms. We package our model into freely available tools, to rapidly investigate consequences of anti-DUX4 therapy.

## Introduction

Facioscapulohumeral muscular dystrophy (FSHD) is a prevalent (^~^12/100,000^1^), incurable, inherited skeletal myopathy. The condition is characterised by progressive fatty replacement and fibrosis of specific muscle groups driving weakness and wasting^2^, which is accelerated by inflammation^3^. FSHD is highly heterogeneous, with both the rate of progression and order of muscle involvement varying dramatically from person to person, even between monozygotic twins^4^. Around 75% of patients exhibit a descending phenotype with weakness beginning in the facial muscles, before progressing to the shoulder girdle and latterly lower limb, while the remaining 25% exhibit a range of ‘atypical’ phenotypes, including facial sparing and lower limb predominant^5^. Despite this clinical range, FSHD is associated with significant morbidity and socioeconomic cost^6^.

Genetically FSHD comprises 2 distinct subtypes: FSHD1 (OMIM: 158900, 95% of cases) and FSHD2 (OMIM: 158901, 5% of cases). Both subtypes bear a unifying epigenetic feature: derepression of the D4Z4 macrosatellite at chromosome 4q35. In FSHD1 this is due to truncation of the D4Z4 macrosatellite from the typical >100 to 10-1 units^7^. In FSHD2, derepression is due to mutation in a chromatin modifier, typically *SMCHD1*^8^ but rarely *DNMT3B*^9^ or *LRIF1*^10^. In addition to D4Z4 epigenetic derepression, FSHD patients also carry certain permissive 4qA haplotypes distal to the last D4Z4 repeat encoding a polyadenylation signal^7^.

Each 3.3kb D4Z4 repeat encodes the transcription factor DUX4, which plays a role in zygotic genome activation, after which it is silenced in somatic tissues^11^. In FSHD, however, epigenetic derepression of the D4Z4 region allows inappropriate transcription of DUX4 from the most distal D4Z4 unit, with transcripts stabilised by splicing to the polyadenylation signal in 4qA haplotypes, allowing translation. Mis-expression of DUX4 protein is thus believed to underlie FSHD pathogenesis and DUX4 inhibition is currently the dominant approach to FSHD therapy^12,13^.

However, DUX4 is extremely difficult to detect in FSHD patient muscle, with the vast majority of transcript and protein level studies failing to detect DUX4 in FSHD muscle biospies^2^. When DUX4 is detected in FSHD patient muscle, it is at very low levels, requiring highly sensitive techniques such as nested RT-qPCR for transcripts^14^ and proximity ligation assays for protein^15^. Investigation of FSHD patient-derived myoblasts have confirmed this very low level of DUX4 expression^2^. Single cell and single nuclear transcriptomic studies find only 0.5-3.8% of in vitro differentiated FSHD myonuclei express DUX4 transcript^16,17^. Immunolabelling studies only detect DUX4 protein in between 0.1-5% of FSHD myonuclei^18,19^.

As DUX4 is a transcription factor, it has been proposed that DUX4 target genes may represent a key driver of FSHD pathology. However multiple meta-analyses have found DUX4 target gene expression to be a poor biomarker of FSHD muscle, except in the context of significant inflammation^20,21^, where it may be confounded by immune cell gene expression^22^. Importantly, a recent phase 2b clinical trial of the DUX4 inhibitor losmapimod failed to reach its primary endpoint of reduced DUX4 target gene expression in patient muscle, despite improvement in functional outcomes^23^.

How can such rare expression of DUX4 drive a pathology as significant as FSHD? DUX4 expression in FSHD patient myoblasts follows a burst-like pattern, and cells expressing DUX4 have significantly shortened lifespans, suggesting a gradual attrition of cells over time^19^. A mouse model in which DUX4 expression is induced in a rare burst-like manner, bears striking histological and transcriptomic similarities to FSHD patient muscle^24,25^. DUX4 expression in FSHD patient derived, multi-nucleated myotubes also displays a gradient across myonuclei, suggesting that DUX4 may ‘diffuse’ either actively or passively from an origin nucleus to neighbouring nuclei, bypassing their need to wait for the rare DUX4 burst^19,26^.

A deeper understanding of DUX4 dynamics and how it drives FSHD pathology is essential to move towards anti-DUX4 therapy. Not only would this explain DUX4 target gene expression as a suboptimal monitoring tool, but would enable optimal therapeutic design, through in silico perturbation of the parameters underlying DUX4 expression and toxicity.

Despite considerable discussion of ‘rare-bursts’ and ‘diffusion’ no attempt has been made to understand DUX4 expression through stochastic processes or differential equations; the natural setting to place these dynamics. Here we combine ordinary differentiation equation models with stochastic gene expression models to construct a tuneable in silico simulation of DUX4 regulation in FSHD cell culture, both in unfused myocytes and syncytial multinucleated myotubes.

Through analysis of iDUX4 myoblasts, scRNAseq and snRNAseq of FSHD differentiated myoblasts we derive experimental estimates for the parameters of our model, including DUX4 mRNA degradation, transcription and translation rates and DUX4 target gene activation rates. Simulation of our model provides a striking fit to DUX4 and DUX4 target gene expressing cell proportions seen in scRNAseq of FSHD myocytes. Importantly, our model predicts that DUX4 drives significant cell death, despite expression limited to <1% of live cells. By comparing scRNAseq of unfused FSHD myonuclei to snRNAseq of multinucleated FSHD myotubes, we find evidence of DUX4 protein syncytial diffusion. We extend our model to examine DUX4 spreading between adjacent myonuclei and project our simulation onto the surface of a muscle fibre, in a spatially relevant model. Employing genetic algorithms to fit our spatial model to snRNAseq of FSHD syncytial myotubes, we provide an estimate for the syncytial diffusion rate of DUX4 protein.

We package our model into three freely available user interfaces, presenting an in silico toolkit to assess the impact of specific anti-DUX4 therapies on FSHD cell culture in a rapid, cost effective and unbiased manner.

## Results

### Compartment and Promoter-switching models of DUX4 expression

Here we consider two models of DUX4 expression in FSHD myocytes, a deterministic model of FSHD cell states we call the compartment model and a stochastic model of gene expression we call the promoter switching model.

We first describe the compartment model. FSHD single myocytes can express DUX4 and therefore DUX4 target genes; DUX4 target gene expression leads to cell death ^19,27,28^. We thus propose that an FSHD single myocyte at a given time *t*, occupies one of the following 5 states/compartments, defined by its transcriptomic distribution:

1. *S*(*t*) – a *susceptible* state where the cell expresses no DUX4 mRNA and no DUX4 target mRNA (DUX4 -ve/Target gene -ve: DUX4 mRNA naive cell)
2. *E*(*t*) – an *exposed* state where the cell expresses DUX4 mRNA but no DUX4 target mRNA (DUX4 +ve/Target gene -ve: DUX4 transcribed but not translated)
3. *I*(*t*) – an *infected* state where the cell expresses both DUX4 mRNA and DUX4 target mRNA (DUX4 +ve/Target gene +ve: DUX4 transcribed and translated)
4. *R*(*t*) – a *resigned* state where the cell expresses no DUX4 mRNA but does express DUX4 target mRNA (DUX4 -ve/Target gene +ve: i.e. a historically DUX4 mRNA expressing cell)
5. *D*(*t*) – a *dead* state

We propose a model in which the single FSHD myocyte can transition through these 5 states according to 5 parameters:

1. *V_D_* – the average transcription rate of DUX4 in a single cell
2. *d*_0_ – the average degradation rate of DUX4 mRNA
3. *T_D_* – the average translation rate of DUX4 from mRNA to active protein
4. *V_T_* – the average transcription rate of DUX4 target genes in the presence of DUX4
5. *D_r_* – the average death rate of DUX4 target positive cells

We allow cells to transition from DUX4 mRNA negative states *S*(*t*) and *R*(*t*) to DUX4 mRNA positive states *E*(*t*) and *I*(*t*) respectively at the rate of transcription of DUX4, *V_D_*, with transition in the reverse direction occurring at the degradation rate of DUX4 mRNA, *d*_0_. Transition from state *E*(*t*) to state *I*(*t*) requires DUX4 mRNA to be translated to active protein and DUX4 target mRNA to be expressed and thus occurs at the rate *T_D_V_T_*. Lastly DUX4 target gene +ve states *I*(*t*) and *R*(*t*) transition to the dead cell state *D*(*t*) at rate *D_r_*. Our compartment model can be represented schematically **(Figure 1A)** or equivalently as a system of ordinary differential equations (ODEs) **(Figure 1B).**

**Figure 1:**
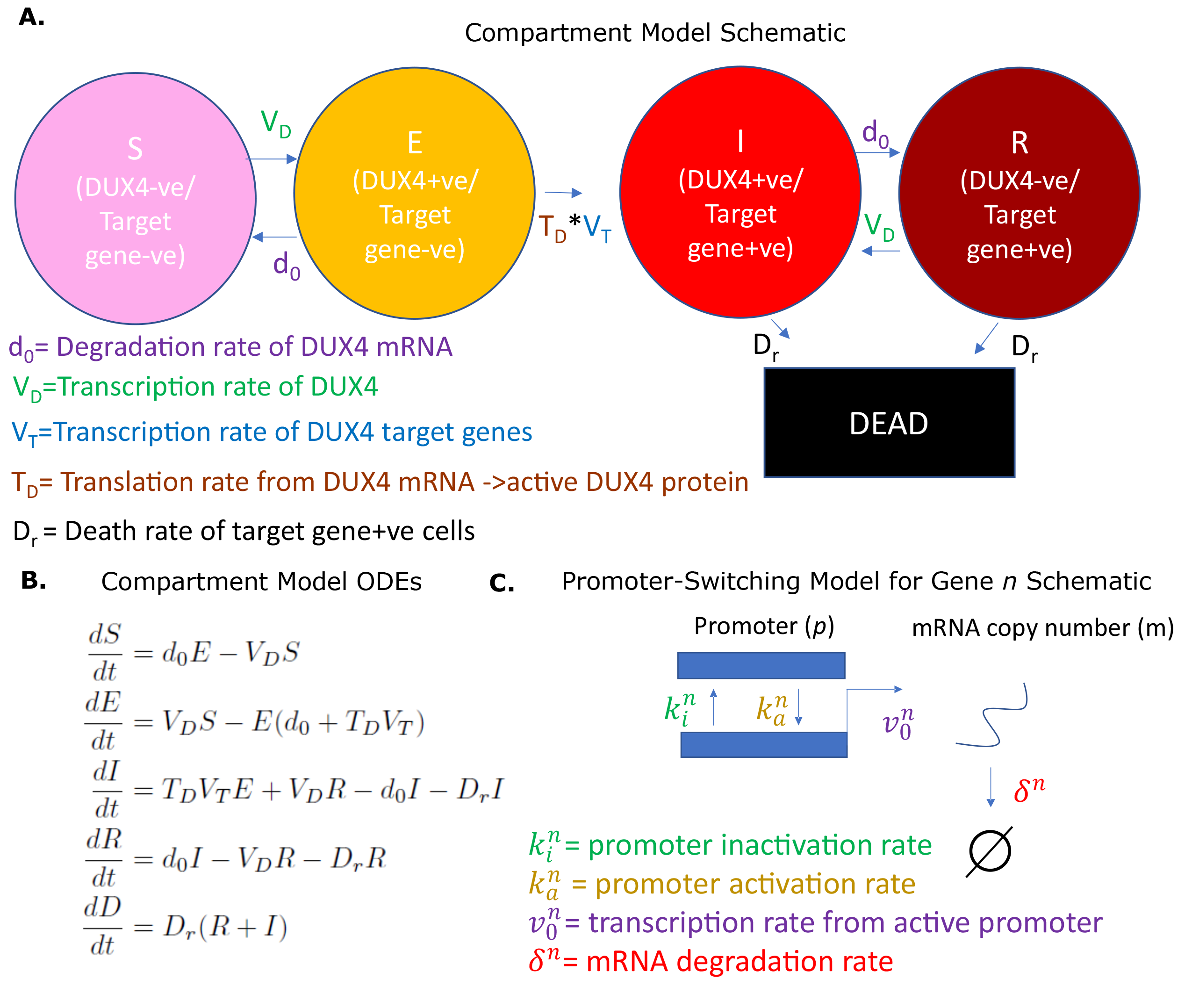
Overview of Models. (A) Schematic of the compartment model describing transition between 5 FSHD states according to 5 underlying parameters. (B) Ordinary differential equations describing the compartment model. (C) Schematic of the promoter-switching model of gene expression.

There are important assumptions in our compartment model:

1. Cells are assumed not to proliferate over the evolution of the model. We restrict applications to differentiating cells which have exited the cell cycle.
2. The death rate of DUX4 target gene -ve cells is negligible in comparison to the death rate of target gene +ve cells. This assumption is justifiable given published data on the death rate of DUX4 target gene positive cells^19^.
3. The transition from DUX4 target gene -ve cell to DUX4 target gene +ve cell states is irreversible, i.e., we assume that the volume of target transcripts induced by DUX4 is sufficiently large, such that the rate of their degradation to zero is negligible in comparison to the death rate of target positive cells. This assumption is justified given that DUX4 is a potent transcriptional activator and pioneer factor^29,30^.

In what follows, we derive experimental estimates for the 5 parameters underlying the compartment model.

A further preliminary is the promoter-switching model. This model is precisely the two-stage telegraph process, which has a long history of study in stochastic gene expression^31–33^. Under this model, we assume that the promoter of a gene *n* can occupy one of two states: active and inactive and that transition from active (*a*) to inactive (*i*) state occurs at a rate 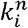, with the reverse transition occurring at rate 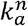. We further assume that an active promoter can transcribe a single mRNA at rate 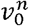, which degrades at rate *δ^n^*. The model is represented schematically in **Figure 1C.** This model of gene expression has been shown to follow a Poisson-Beta distribution^32,33^, where the promoter state is determined by a Beta-distributed variable 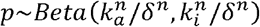, and the mRNA copy number distribution conditional on the promoter state follows a Poisson distribution 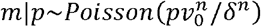. Under this interpretation maximum likelihood estimates (MLEs) for the normalised underlying parameters 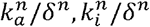 and 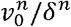 can be approximated from single cell transcriptomic data^32,33^.

We assume that the proportion of time the promoter is in the active state, multiplied by the transcription rate of the active promoter in our promoter-switching model 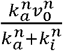, is a reasonable proxy for the average rate of transcription of a gene in our compartmental model. We employ this assumption to estimate the average transcription rates *V_D_* and *V_T_* for the compartment model.

### Estimating the kinetics of DUX4 mRNA

Our compartment model contains 2 parameters governing the kinetics of DUX4 mRNA: the degradation rate *d*_0_ and the average transcription rate *V_D_*.

To estimate the degradation rate *d*_0_ we employed human immortalised LHCN-M2 myoblasts expressing DUX4 under the control of a doxycycline inducible promotor (iDUX4 myoblasts)^29^. DUX4 expression was induced with 250 ng/ml doxycycline for 7 hours, a level and duration we have found sufficient to induce robust DUX4 mRNA expression, but only weak activation of DUX4 target genes and no widespread apoptosis^34^. After 7 hours of induction, myoblasts were washed and supplemented with fresh medium without doxycycline. Samples were harvested for RNA extraction in triplicate immediately after washing and then at 1, 2, 3, 4, 5, 6, 8 and 10 hours post wash. RT-qPCR was performed employing a standard curve to quantify DUX4 mRNA copy number over our time course, to monitor DUX4 mRNA degradation in the absence of induction **(Figure 2A).**

**Figure 2:**
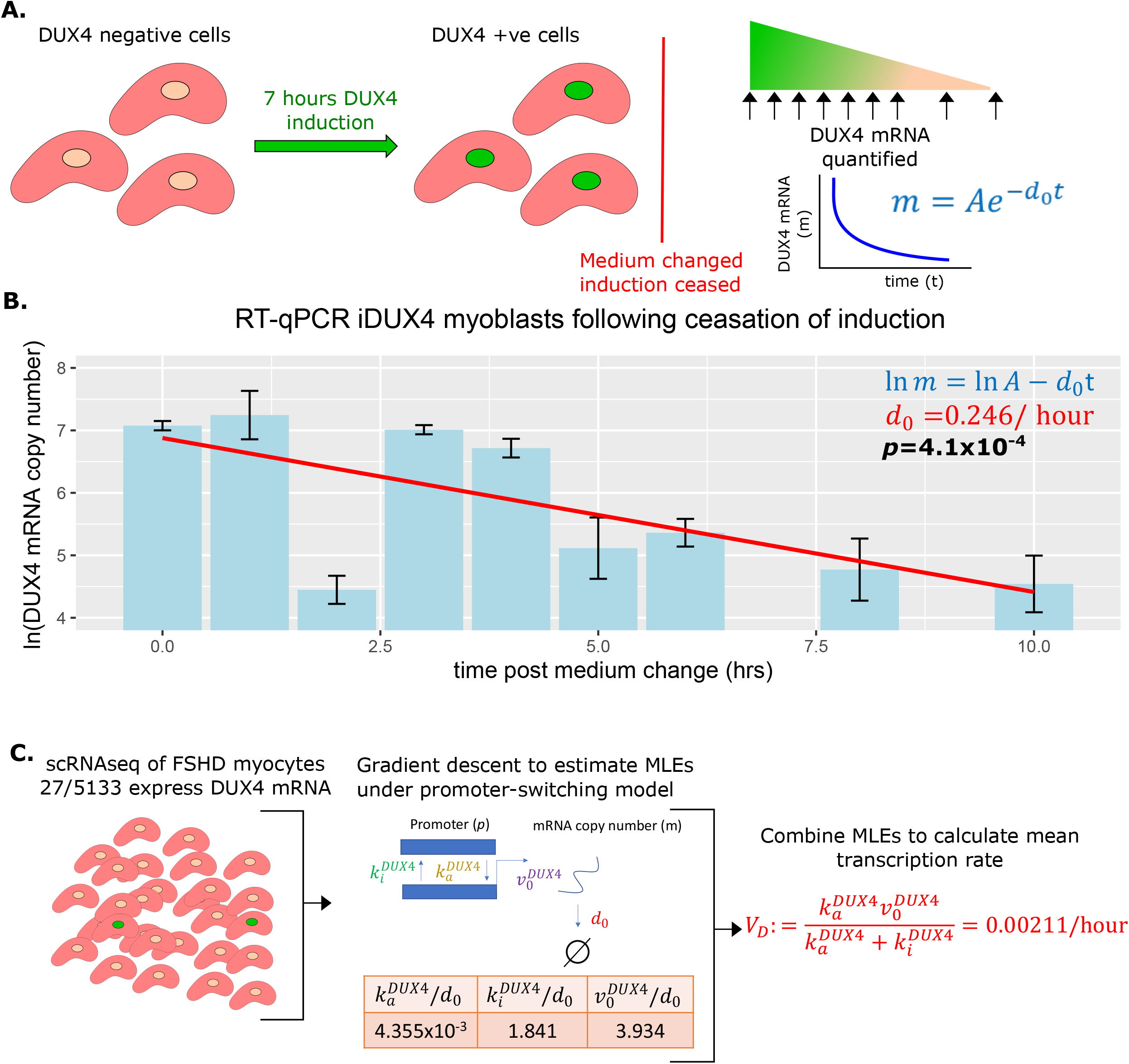
Estimation of DUX4 mRNA degradation rate (d_0_) and average transcription rate (V_D_). (A) Schematic of experiment performed to estimate the DUX4 mRNA degradation rate *d*_0_. iDUX4 myoblasts were induced to express DUX4 with 250 ng/ml of doxycycline for 7 hours before doxycycline was washed away. DUX4 mRNA was quantified at multiple time points post wash using RT-qPCR. (B) Bar chart displays RT-qPCR of ln(DUX4 copy number) at post-wash times 0, 1, 2, 3, 4, 5, 6, 8 and 10 hours. Bar represents the average of triplicates and standard error of the mean is displayed. A line of best fit of ln(DUX4 copy number) against time is displayed alongside the corresponding linear regression *p*-value (bold) and the slope of the line corresponds to *d*_0_ (red). (C) Schematic of the estimation of average DUX4 transcription rate *V_D_* from scRNAseq data of 5133 FSHD myocytes. The MLEs for the underlying parameters of DUX4 transcription under the promoter-switching model are estimated via gradient descent and combined to estimate the average DUX4 transcription rate (red text).

As anticipated DUX4 mRNA levels decayed exponentially over time in the absence of doxycycline (linear regression of ln(DUX4 mRNA) vs time, *p*=4.1×10^-4^, **Figure 2B),** allowing us to calculate the degradation rate of DUX4 mRNA, *d*_0_ = 0.246/hour. This estimate suggests that the half life of DUX4 mRNA is approximately 2.8 hours, not atypical for a transcription factor, and on the faster end of the mRNA degradation distribution^35^.

To estimate the mean transcription rate of DUX4, *V_D_*, we applied the promoter-switching model presented above. We considered the scRNAseq dataset of FSHD single myocytes produced by van den Heuvel et al., 2019^17^. This dataset comprises 7047 single myocytes, which were differentiated for 3 days in the presence of EGTA to inhibit fusion. 5133/7047 (73%) single myocytes were derived from 4 FSHD patients (2 FSHD1 and 2 FSHD2). DUX4 mRNA expression was detected in 27/5133 (0.53%) single FSHD myocytes but not in control myocytes^17,21^. Considering the 5133 FSHD myocytes together we implemented a gradient descent algorithm to approximate the MLEs of the normalised parameters 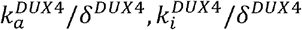 and 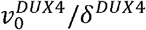 from the Poisson-Beta interpretation of the promoter-switching model of DUX4 expression **(Figure 2C)**.

We note that *δ^DUX4^* = *d*_0_, which we have already estimated, this enabled us to renormalise these parameters and compute the average DUX4 transcription rate, 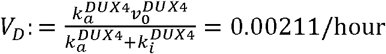.

### Estimating kinetics of DUX4 target activation

We next estimated the average transcription rate of the DUX4 target genes in the presence of DUX4 mRNA, *V_T_*. We focused on 8 DUX4 target genes: *ZSCAN4, TRIM43, RFPL1, RFPL2, RFPL4B, PRAMEF1, PRAMEF2* and *PRAMEF12* that have been identified as direct DUX4 targets via ChIP-seq^36^. We have shown that these 8 genes are the only features consistently up-regulated in human myoblasts expressing DUX4, across 4 independent studies^2,19,29,37,38^.

Examining mRNA levels of these 8 DUX4 target genes in scRNAseq^17^ and snRNAseq^16^ studies of FSHD and control differentiated myoblasts, expression was restricted to FSHD cells/nuclei and never observed in controls. This pattern of expression mirrors that of DUX4 mRNA and suggests that these targets are highly specific and are unlikely to be activated in the absence of DUX4.

To confirm our hypothesis, we applied the promoter-switching model. We returned to the scRNAseq data of 5133 FSHD myocytes generated by van den Heuvel et al., 2019^17^ and divided the data into the 27 DUX4 +ve cells and 5106 DUX4 -ve cells. For each of our 8 DUX4 target genes, we implemented our gradient descent algorithm to compute the MLEs of the normalised parameters 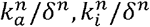 and 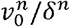, underlying a promoter-switching model for the given gene across the 27 DUX4 +ve FSHD myocytes and the 5106 DUX4 -ve FSHD myocytes separately **(Figure 3A).**

**Figure 3:**
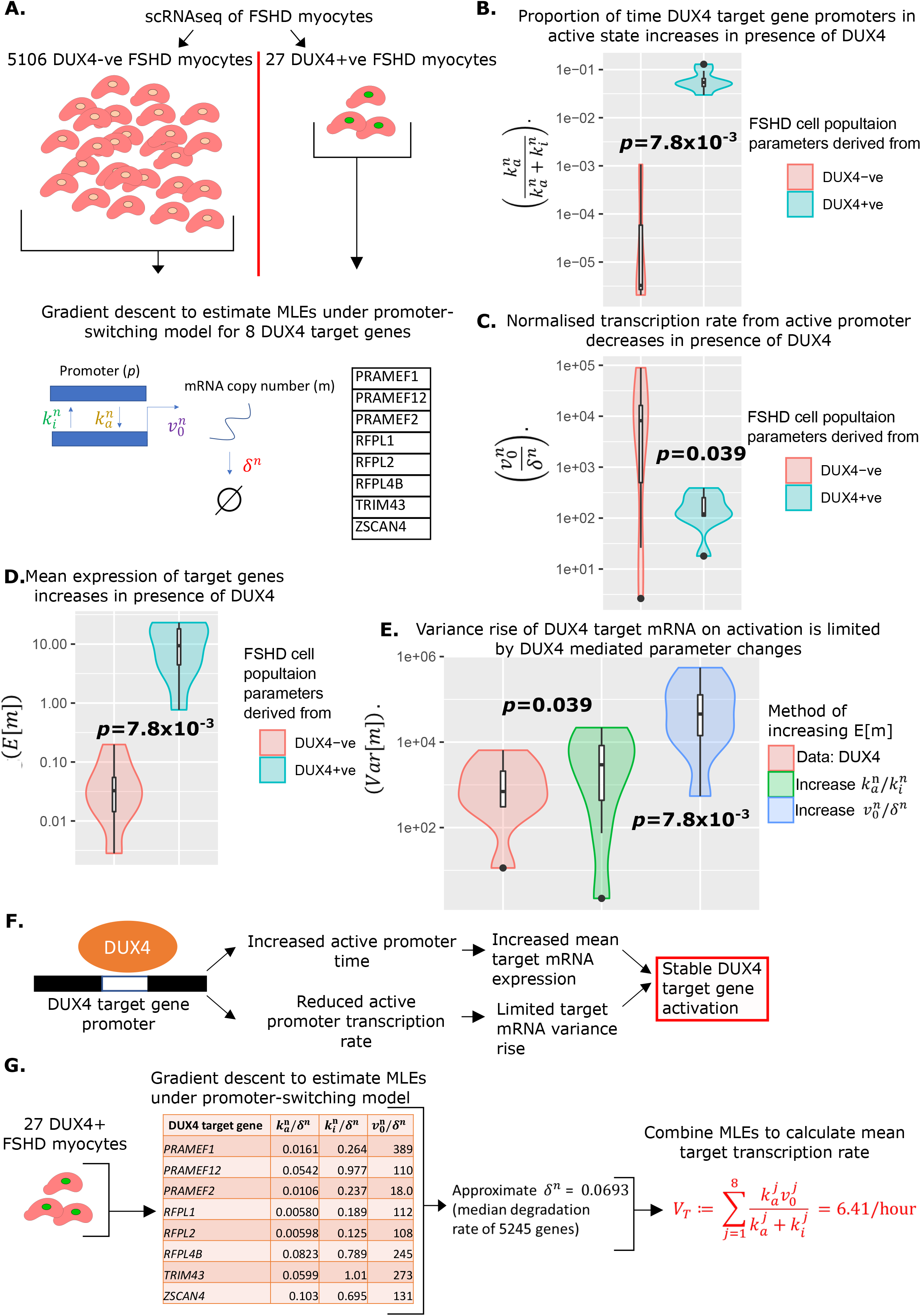
Estimation of promoter-switching model parameters for DUX4 target genes and estimation of V_T_. (A) Schematic of estimation of promoter-switching model parameters for 8 DUX4 target genes, across 5106 DUX4 -ve FSHD single myocytes and 27 DUX4 +ve FSHD single myocytes from van den Heuvel et al., 2019^17^. Violin plots display (B) proportion of time in active promoter state (C) normalised active promoter transcription rate and (D) mean mRNA copy number, for the 8 DUX4 target genes in the 5106 DUX4 -ve and 27 DUX4 +ve FSHD myocytes separately, *p*-values correspond to 2 tailed paired Wilcoxon tests. (E) Violin plot displays the variance of mRNA copy number for the 8 DUX4 target genes calculated from the 27 DUX4 +ve FSHD myocytes (red) and calculated assuming DUX4 up-regulates targets only via increase in (blue) the normalised transcription rate 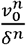 or (green) the ratio of active to inactive promoter transition rates 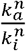. Paired two-tailed Wilcoxon *p*-values are displayed comparing adjacent distributions. (F) Schematic displaying how the change in parameters underlying the promoter-switching models for the 8 DUX4 target genes in the presence of DUX4 leads to stable target gene activation. (G) Schematic of the estimation of the average DUX4 target gene transcription rate *V_T_*, from scRNAseq data of 27 DUX4 +ve FSHD myocytes.

In the presence of DUX4 mRNA the proportion of time the promoters of the 8 DUX4 target genes remained in the active state, 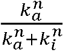, significantly increased (paired Wilcoxon *p*=7.8×10, **Figure 3B),** as expected. Curiously however, the normalised rate of transcription of the DUX4 target genes from the active promoter, 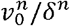, significantly decreased in the presence of DUX4 (paired Wilcoxon *p*=0.039, **Figure 3C).**

To investigate further, we considered the moments of the distribution of mRNA copy number, *m*, under the promoter-switching model. It can be shown that the mean mRNA copy number satisfies^32^:

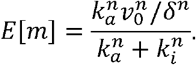

The changes in parameters we calculate for the DUX4 target genes confirmed that in the presence of DUX4 mRNA, the mean expression of all 8 DUX4 targets increases (paired Wilcoxon *p*=0.039, **Figure 3D),** i.e., the drop in 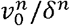 is over-compensated for by the rise in 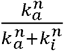. However, it is curious that 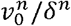 should drop at all.

It can be shown that the variance of mRNA copy number *m*, satisfies^32^:

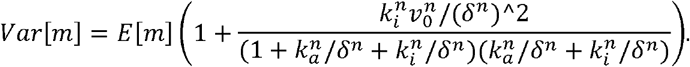

This expression ensures that as the mean mRNA copy number rises, so too must the variance, however the level to which the variance rises is controlled by a term that is monotonic increasing in 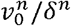 and depends on the promoter state parameters in a more complex way.

We postulated the pattern of DUX4 target gene parameter changes we observe in the presence of DUX4, have the effect of raising mean target mRNA expression, while suppressing target mRNA variance, i.e., a controlled activation of target genes. To investigate this, we considered 2 hypothetical scenarios in which the mean expression of target mRNAs, *E*[*m*] in the absence of DUX4, is increased to the same level we observe in the presence of DUX4. In the first scenario we considered only increasing 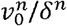 to achieve the rise in *E*[*m*], in the second we considered only increasing the ratio of active to inactive promotor 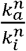. Both hypothetical scenarios resulted in a significantly higher variance of target mRNA than observed in our data, with the pure rise in 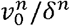 driving the most dramatic increase in target mRNA variance **(Figure 3E).**

Taken together these results suggest that under our promoter-switching model, increasing the expression of a gene comes at the cost of increasing the variance of its expression, and that the greatest contributor to this variance comes from the normalised active promoter transcription rate 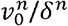. The parameter changes we observe in DUX4 target genes, suggest a management of this trade off by DUX4, which increases the mean expression of target mRNA through a large increase in the proportion of time the promoter is active, 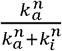, while offsetting the resulting rise in variance through a modest decrease in normalised active promotor transcription 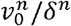 **(Figure 3F).**

We formulate the mean transcription rate of at least one of the 8 DUX4 targets from our compartment model, *V_T_*, as the sum of the mean transcription rates of all 8 target genes in the presence of DUX4 mRNA, i.e.:

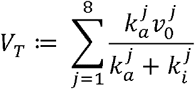

where *j* indexes the 8 DUX4 target genes, and the promoter-switching model parameters are estimated from the 27 DUX4 expressing FSHD single myocytes. Our promoter-switching model scRNAseq derived MLEs are normalised parameters 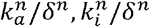 and 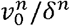. We must therefore estimate *δ^n^* for each target gene to compute *V_T_*. As there are a range of target genes, we approximate *δ^n^* for each by the median mRNA degradation rate observed in an analysis of 5245 genes^35^, and set *δ^n^* = 0.0693 resulting in *V_τ_* = 6.41/hour **(Figure 3G).**

The remaining 2 parameters of our compartment model were calculated from published data. For the translation rate of DUX4 mRNA to active protein *T_D_*, we considered our analysis of iDUX4 myoblasts^34^. We induced DUX4 expression with 250 ng/ml doxycycline and performed RT-qPCR to assess expression of *DUX4*, and 3/8 of our DUX4 target genes *ZSCAN4, TRIM43* and *PRAMEF1*, at 7 hours, 16 hours and 24 hours of induction. DUX4 mRNA levels peaked at 7 hours, while the expression of DUX4 target genes peaked between 16 and 24 hours^34^. This suggests a delay between DUX4 mRNA production and the presence of active DUX4 protein of between 9 and 17 hours, on average 13 hours. We thus estimate 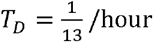.

For the death rate of DUX4 target positive cells, *D_r_*, we consider the data of Rickard et al., 2015^19^, in which differentiating FSHD myoblasts containing a DUX4 activated GFP reporter were imaged every 15 mins for 120 hours. Following activation of the DUX4 reporter, cells died ^~^20.2 hours later^19^. We thus estimate 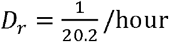.

### Compartment model simulation

Having defined experimental estimates for parameters underlying the compartment model, we simulated the model forward in time to observe how an initial distribution of cells progresses through the 5 compartments.

To provide a ground truth we considered the scRNAseq data of van den Heuvel et al., 2019^17^. Defining a DUX4 target gene +ve cell as expressing at least one of our 8 DUX4 target genes, we assign the 5133 FSHD myocytes to one of the four live cell states of our compartment model: *S*(3 *days*) = 4956 (96.6%), *E*(3 *days*) = 14 (0.273%), *I*(3 *days*) = 13 (0.253%), *R*(3 *days*) = 150 (2.92%).

If we assume that at the start of differentiation all FSHD myocytes occupy state *S*(0) being DUX4 negative and DUX4 target gene negative, simulating our model we estimated that 7% of starting cells will have died over 3 days. To replicate the starting conditions of the scRNAseq data we thus set *S*(0) = 5133(1 + 0.07) = 5488, *E*(0) = *I*(0) = *R*(0) = *D*(0) = 0. Simulating our model over 3 days from this starting condition, we obtained cell state proportions statistically indistinguishable from the experimental scRNAseq data: *S*(3 *days*) = 4953 (96.5%), *E*(3 *days*) = 14 (0.253%), *I*(3 *days*) = 25 (0.487%), *R*(3 *days*) = 116 (2.26%) (Chi-Squared p=0.99, **Figure 4A).**

**Figure 4:**
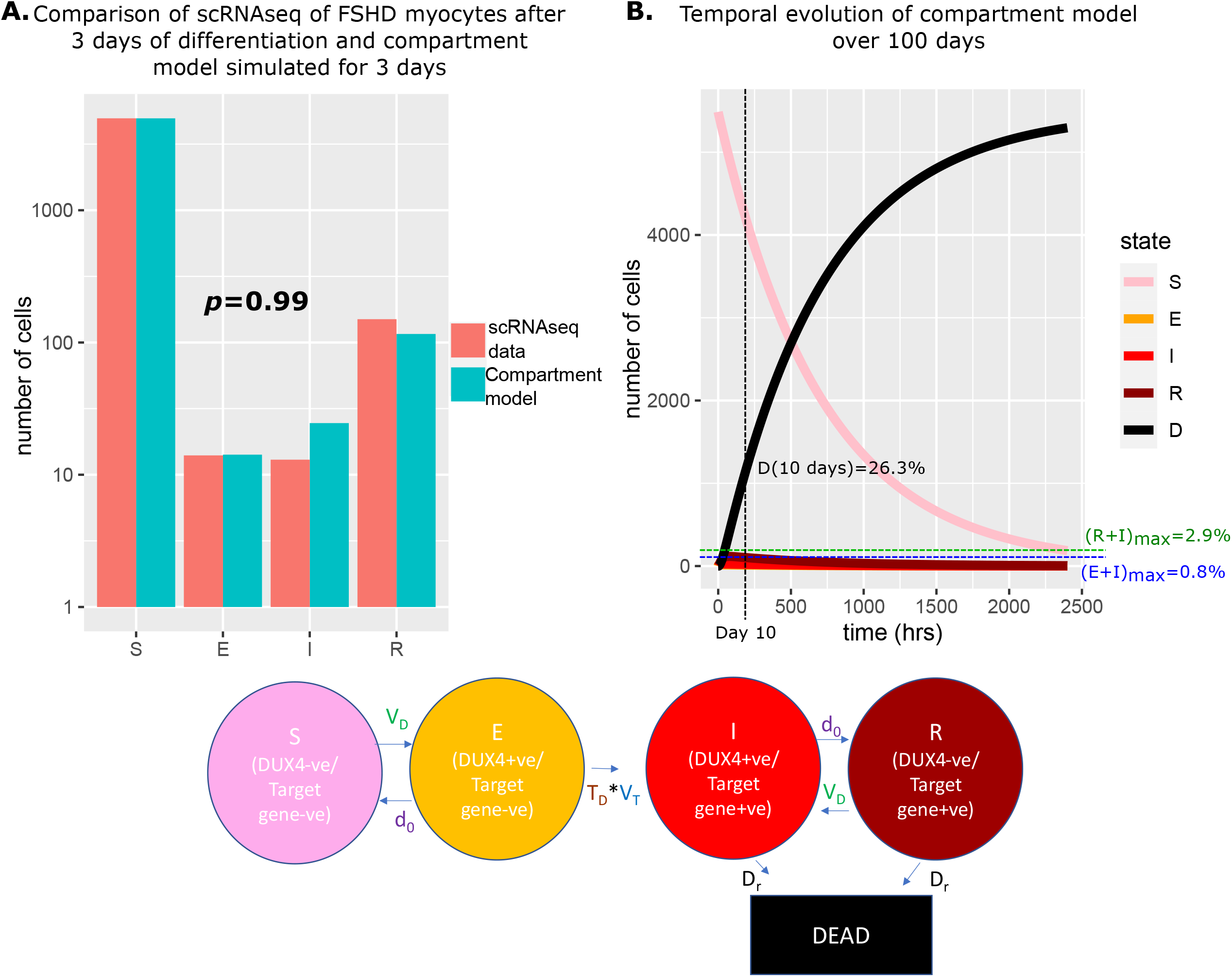
Simulation of the compartment model and comparison with scRNAseq data of unfused FSHD myocytes. (A) Bar chart displays the number of cells in each live cell compartment (blue) in our model following simulation over 3 days using experimentally estimated parameters and (red) in 5133 single FSHD myocytes. A Chi-squared goodness of fit *p*-value tests the alternate hypothesis that the two distributions are different. (B) Line plot displaying how the number of cells in each of the 5 compartments of the model changes over 100 days from a starting state of 5133(1+0.07) cells. The percentage of cells dead after 10 days is displayed alongside the maximum percentage of live cells which are DUX4 +ve (E+I)_max_ and DUX4 target gene +ve (R+I)_max_.

We next simulated our model forward over 100 days to observe how the proportion of cells in each state changed **(Figure 4B).** As expected, the number of cells in the DUX4 naive, susceptible state *S*(*t*) gradually decreased, while the number of dead cells *D*(*t*) gradually rose, so that after 10 days 26.3% of cells had died. Remarkably despite cell death in our model only being attributable to DUX4 expression, the dynamics predict that this is achieved while keeping the number of DUX4 mRNA and DUX4 target mRNA positive cells extremely low. DUX4 positive cells never rose to more than 0.8% of the live cell population and DUX4 target positive cells never more than 2.9%.

Our compartment model with experimentally derived parameters thus provides an excellent fit to real world data and gives mechanism to how extremely low levels of DUX4 and target gene expression can drive significant cell death in FSHD myocytes.

### Modelling DUX4 expression in FSHD myotubes allowing syncytial diffusion

Having considered scRNAseq data of unfused FSHD myocytes differentiated for 3 days, in which DUX4 expressed in a given nucleus cannot interact with other nuclei, we next considered syncytial diffusion of DUX4. Jiang et al., 2020^16^ published single myonuclear RNAseq (snRNAseq) of FSHD2 myoblasts differentiated over 3 days to form multinucleated myotubes. The data describes 139 FSHD2 myonuclei and 76 control myonuclei. As with the scRNAseq data^17^, none of the control myonuclei express DUX4 nor any of the 8 DUX4 target genes. For FSHD myonuclei, 3/139 (2.2%) express DUX4. The number of myonuclei in each of the live ‘cell’ compartments of our model are: *S*(3 *days*) = 58 (41.7%), *E*(3 *days*) = 0 (0%), *I*(3 *days*) = 3 (2.2%), *R*(3 *days*) = 78 (56.1%).

Comparing the proportions of myonuclei (cells) assigned to the 4 live states of our compartment model in the unfused FSHD scRNAseq and syncytial FSHD snRNAseq data sets, we found significant differences in the distributions (Chi-Squared *p*<2.2×10^-16^, **Figure 5A).** The unfused FSHD myocytes demonstrated a higher proportion of DUX4 -ve, DUX4 target gene -ve *susceptible (S)* cells, while the syncytial FSHD myonuclei demonstrated a greater proportion of DUX4 -ve, DUX4 target gene +ve *resigned (R)* cells.

**Figure 5:**
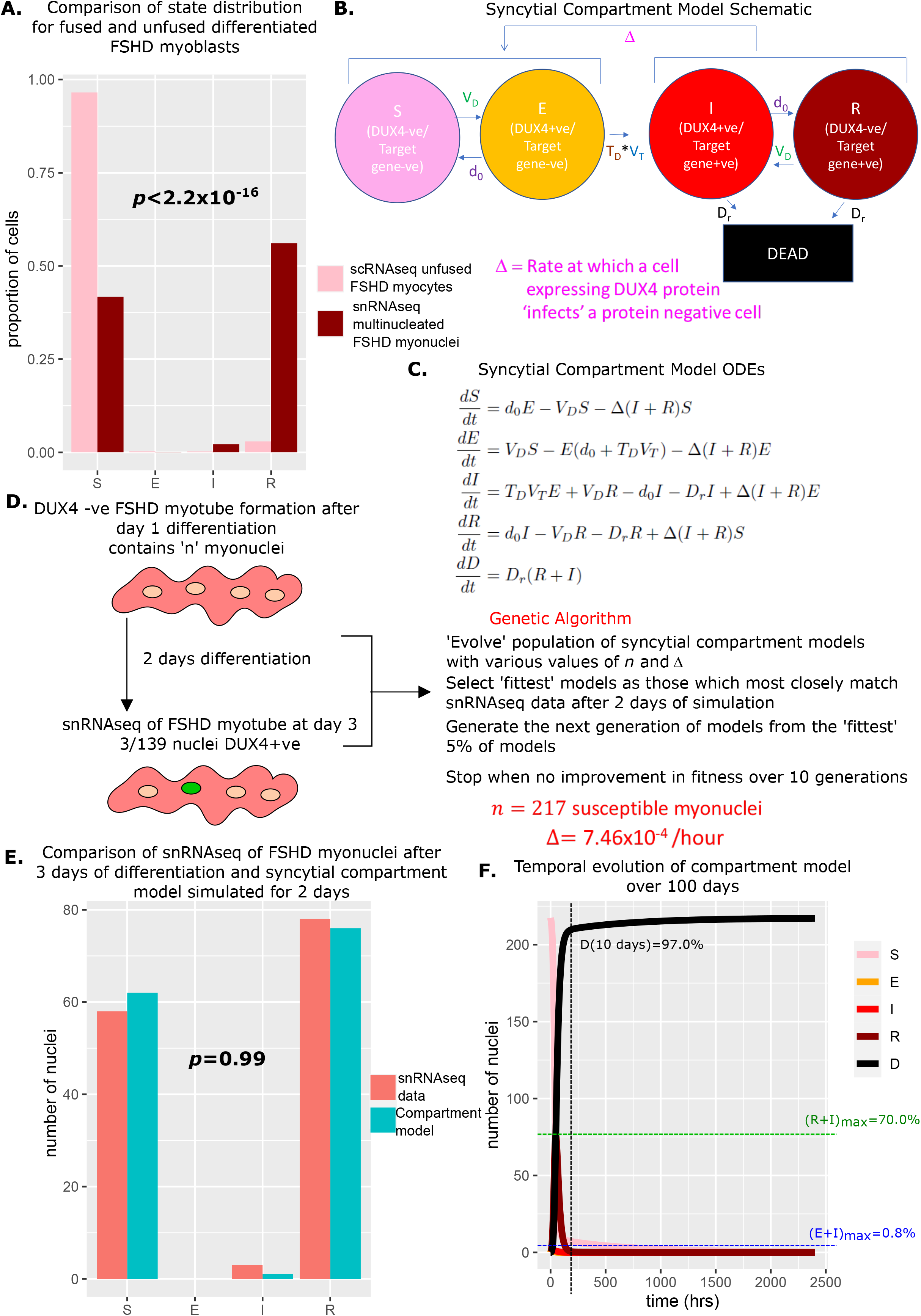
Derivation and simulation of the syncytial compartment model and comparison with snRNAseq data of fused FSHD myonuclei. (A) Bar chart displays the proportion of cells in each live compartment of our model in (pink) 5133 unfused FSHD single myocytes from van den Heuvel et al.,^40^ and (dark red) 139 fused FSHD single myonuclei of Jiang et al.^16^. A Chi-squared goodness of fit *p*-value tests the alternate hypothesis that the two distributions are different. (B) Schematic of the syncytial compartment model, allowing DUX4 protein permissive states *I* and *R*, to ‘infect’ non-permissive states *S* and *E*. (C) Ordinary differential equations describing the syncytial compartment model. (D) Schematic of the genetic algorithm employed to estimate the DUX4 diffusion rate Δ and the starting population size *n*, for the syncytial compartment model from 139 fused FSHD single myonuclei. (E) Bar chart displays the number of cells in each live cell compartment (blue) in our syncytial compartment model following simulation over 2 days using experimentally estimated parameters and (red) in 139 fused single FSHD myonuclei. A Chi-squared goodness of fit *p*-value tests the alternate hypothesis that the two distributions are different. (F) Line plot displaying how the number of cells in each of the 5 compartments of the syncytial compartment model changes over 100 days from a starting state of *n* cells. The percentage of cells dead after 10 days is displayed alongside the maximum percentage of live cells which are DUX4 +ve (E+I)_max_ and DUX4 target gene +ve (R+I)_max_.

We investigated whether allowing diffusion of DUX4 protein between myonuclei in our model could explain this difference in proportions. Two states in our compartment model express DUX4 target genes and thus have evidence for DUX4 protein: *I*(*t*) and *R*(*t*), while states *S*(*t*) and *E*(*t*) do not. We updated our model to the syncytial compartment model, where we allow the DUX4 protein compatible states to ‘infect’ the DUX4 protein incompatible states at a rate Δ **(Figure 5B&C).**

We next employed a genetic algorithm to fit our syncytial compartment model to the snRNAseq data^16^. As differentiating FSHD myoblasts take approximately 24 hours to initiate fusion^39^, and DUX4 expressing cells show defects in fusion^30^, we assumed that day 3 of differentiation represents day 2 of syncytial myonuclei and simulated our model over 48 hours, from a starting state of *S*(0) = *n, E*(0) = *I*(0) = *R*(0) = 0. We optimised 2 parameters via our genetic algorithm: the DUX4 syncytial diffusion rate Δ, and the number of starting myonuclei in the susceptible state *n* **(Figure 5D).**

The algorithm converged on a solution of *n* = 217 susceptible myonuclei and Δ=7.46×10^-4^/hour. Simulating the syncytial compartment model over 2 days employing these parameters resulted in a statistically indistinguishable approximation to the snRNAseq data^16^: *S*(3 *days*) = 62 (44.6%), *E*(3 *days*) = 0 (0%), *I*(3 *days*) = 1 (0.720%), *R*(3 *days*) = 76 (54.7%), (Chi-Squared *p*=0.99, **Figure 5E).** Thus differences in proportions of cell states in fused versus unfused myocytes can be explained by addition of a diffusion term for DUX4 protein, within the limits of the assumptions of our model.

Simulating the syncytial compartment model over 100 days, we found a much faster death rate than the unfused myocyte model, with 97% of myonuclei dead by day 10. DUX4 mRNA expressing cells remain low in proportion, never exceeding 0.8% of total live nuclei, however DUX4 target gene mRNA expressing cells can now comprise a significant proportion of living nuclei, up to 70% at 73 hours, though on average comprise only 5.6% of living nuclei **(Figure 5F).**

### A cellular automaton model of FSHD myotubes

A limitation of our syncytial compartment model is the assumption that any DUX4 protein positive myonucleus can ‘infect’ any DUX4 protein negative myonucleus. In practice, DUX4 can likely only spread between adjacent myonuclei in a short range interaction^19,26^. To overcome this limitation, we re-cast our compartment model as a cellular automaton.

In this grid based model, squares on the grid represent myonuclei, by introducing an appropriate boundary condition, the grid is topologically equivalent to the surface of a cylinder and thus can be considered to represent myonuclei residing on the surface of a myofiber **(Figure 6A).**

**Figure 6:**
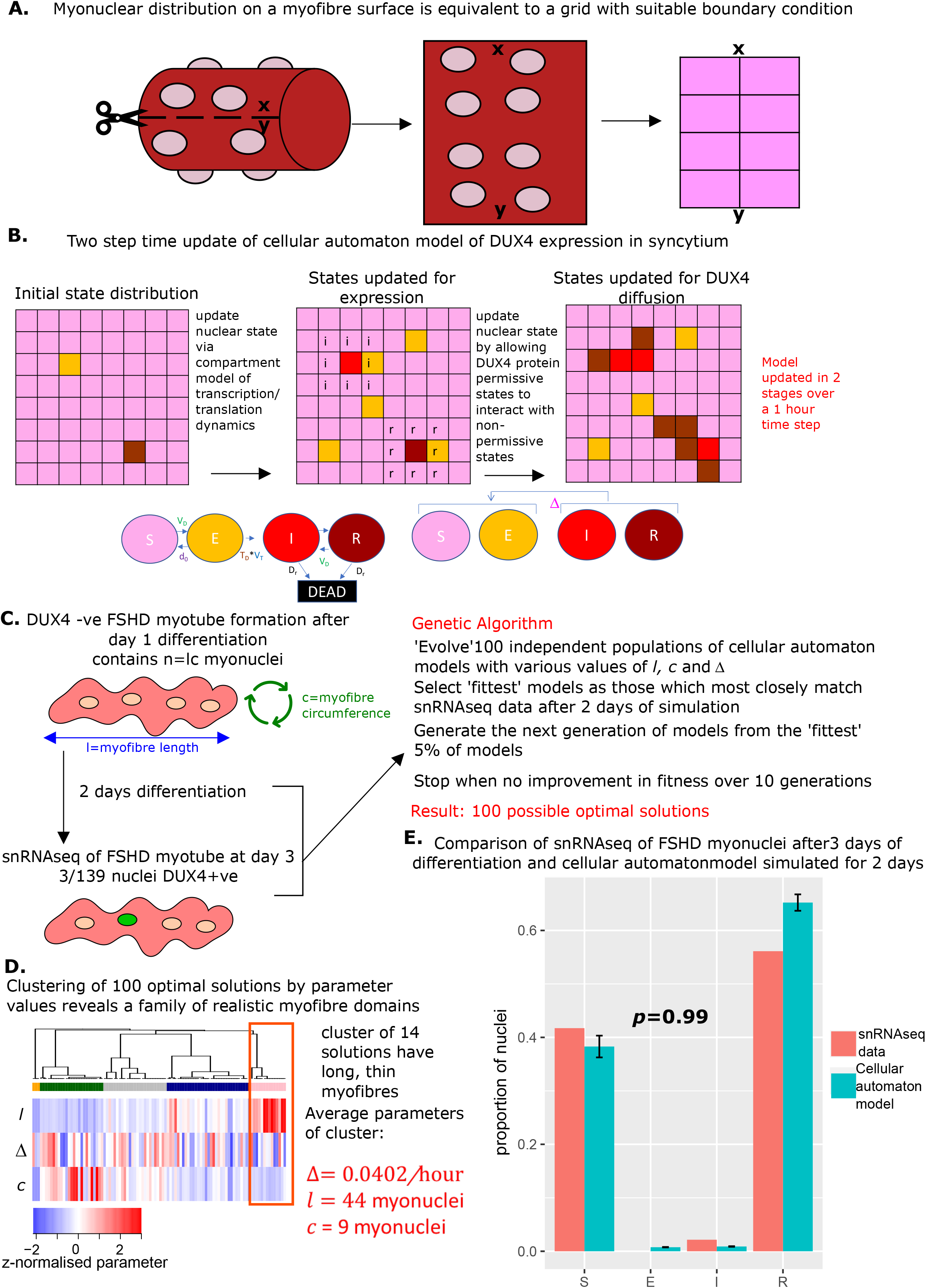
Derivation and simulation of the cellular automaton model and comparison with snRNAseq data of fused FSHD myonuclei. (A) Schematic demonstrating how with suitable boundary condition a grid can be topologically equivalent to a cylinder, and thus encapsulate dynamics on the surface of a myofibre. (B) Schematic of the cellular automaton model of DUX4 expression in a syncytium with grid squares representing single myonuclei updating in 1 hour time-steps via a 2 stage process. (C) Schematic of the genetic algorithm employed to estimate the DUX4 diffusion rate Δ, myotube length *l* and myotube circumference *c* for the cellular automaton model from 139 fused FSHD single myonuclei. (D) Heatmap displays the clustering solution of the 100 optimal parameter regimes *l, c* and Δ produced by fitting 100 genetic algorithms to 139 fused FSHD myonuclei. The highlighted cluster represents a parameter regime in which myotubes are long and thin, the average parameter values of this cluster are displayed in red. (E) Bar chart displays the average proportion of cells in each live cell compartment (blue) in our cellular automaton model following simulation over 2 days using experimentally estimated parameters, alongside standard error of the mean, and (red) in 139 fused single FSHD myonuclei. A Chi-squared goodness of fit *p*-value tests the alternate hypothesis that the two distributions are different.

We evolved our cellular automaton forwards in time in two steps. First, the non-syncytial compartment model was applied to each myonucleus to update the internal state of each nucleus stochastically according to the experimentally derived transition rates and the time period that has elapsed. In the second step, myonuclei in DUX4 protein compatible states *I*(*t*) and *R*(*t*) can ‘infect’ any of the 8 neighbouring myonuclei in the grid which are in states *S*(*t*) or *E*(*t*) at a stochastic rate Δ **(Figure 6B).** Our cellular automaton model, thus retains all features of the compartment model, but facilitates a more realistic topology for a muscle fibre and prevents DUX4 protein from engaging in long-range non-physiological interactions.

We next employed a genetic algorithm to fit the cellular automaton to the syncytial snRNAseq data^16^. We again assumed that day 3 of differentiation corresponds to day 2 of syncytial myotubes and simulate our model over 48 hours, in time steps of 1 hour, from a starting state of *S*(0) = *n, E*(0) = *I*(0) = *R*(0) = 0. However, we now optimised 3 parameters: the DUX4 local diffusion rate Δ, the myotube nuclear length *l* and the myotube nuclear circumference *c*, where *n* = *lc*.

As the cellular automaton model is stochastic due to the discrete time steps employed, one cannot expect the same set of parameters to yield the same distribution of cell states after 48 hours of simulation. Thus the solution of a genetic algorithm may represent a parameter regime optimal for replicating the snRNAseq data, or may represent an unlikely realisation of a parameter regime sub-optimal for replicating the data. To overcome this limitation we implemented 100 genetic algorithms to obtain a ‘family’ of parameter values Δ, *l* and *c*, which underlie potentially optimal solutions **(Figure 6C).**

We performed a clustering analysis of the 100 optimal solutions, employing the parameter values Δ, *l* and *c* as a feature space. This revealed that a cluster of 14 solutions has significantly greater myotube length than circumference (Wilcoxon *p*=5.0×10^-8^). This implies that for these 14 solutions the genetic algorithm has converged to the general structure of a muscle fibre cylinder (long and thin). We focused on the mean parameter values of this cluster as the most physiologically realistic solution: Δ= 0.0402/hour, *l* = 44 myonuclei and *c* = 9 myonuclei **(Figure 6D).**

Simulating the cellular automaton model 100 times with this parameter range over 48 hours resulted on average in a statistically indistinguishable approximation of the snRNAseq syncytial FSHD myonuclei state distribution (Chi-squared *p*=0.98, **Figure 6E).**

Thus after restricting DUX4 protein spreading to only neighbouring nuclei, we were still able to explain the difference between unfused and fused cell state distributions purely with a diffusion term for DUX4 protein.

### A toolbox for assessing the impact of therapies on DUX4 mediated myotoxicity

Anti-DUX4 therapy for FSHD can target any aspect of DUX4 expression. To understand the impact of a given therapeutic strategy one requires two key pieces of information:

1. The true value of the parameters underlying DUX4 expression.
2. How a proposed change in these parameters by a therapeutic will impact cell death.

Here we have derived experimental estimates for the parameters underlying DUX4 expression which can be modified by anti-DUX4 therapy, while our compartmental and cellular automaton models provide the framework for investigating how parameter changes will impact cell death.

To facilitate other investigators using our models to guide anti-DUX4 therapy development, we have packaged them into graphical user interfaces to provide 3 user friendly tools.

The first tool allows investigators to visualise the compartment model and syncytial version, for a range of parameter choices. Sliders allow the investigator to perturb the 6 parameters of our models: *V_D_*, *d*_0_, *V_T_, T_D_, D_r_* and Δ, and outputs the temporal evolution of the models over as many hours as required, from any starting number of *susceptible* cells/nuclei, as well as histograms comparing the perturbed model to the experimental data of van den Heuvel et al.,^40^ and Jiang et al.^16^.

The second tool implements the cellular automaton model of DUX4 in a syncytium. As with the first tool users can employ sliders to perturb the 6 parameters of the model *V_D_, d*_0_, *V_T_, T_D_, D_r_* and Δ, as well as the dimensions of the myofiber grid *l* and *c*, and the number of hours over which the automaton should be simulated. The automaton can then be started and the nuclear state updates will be dynamically played on a grid, while a histogram dynamically updates proportions of nuclei in each state.

The final tool allows investigators to compare dead cell proportions over time for a chosen parameter regime (*V_D_, d*_0_, *V_T_, T_D_, D_r_* and Δ, *l* and *c*), with cell death under our experimentally derived parameters. Users can again select their parameter regime and dynamically view changes in dead cell proportions under the single cell compartment model, to simulate realisations of the cellular automaton model and perform comparisons of cell death via Cox proportional hazard models.

The tools are hosted at the following web addresses:

1. Compartment Models: https://crsbanerji.shinyapps.io/compartment_models/
2. Cellular Automaton: https://crsbanerji.shinyapps.io/ca_shiny/
3. Survival Analysis: https://crsbanerji.shinyapps.io/survival_sim/

The three tools can be used to understand how anti-DUX4 therapies aimed at different model parameters alter the proportion of cells in each compartment of our model at given times. This can guide the optimal therapeutic approaches, as well as optimal time points to assay to validate given therapies using cell culture approaches.

## Discussion

Anti-DUX4 therapy is the leading candidate for an FSHD treatment, with several compounds currently in clinical trials^12,23,41^. However, DUX4 expression in FSHD muscle demonstrates a complex dynamic with DUX4 mRNA, protein and target gene accumulation all difficult to detect^2,21^. Understanding this complex dynamic is essential to the construction of optimal therapy, as it is currently unclear which stage of the DUX4 ‘central-dogma’ one should target to have the most significant impact on pathology. This has led to diverse strategies, targeting epigenetic regulation of DUX4^8,42^, DUX4 mRNA^43,44^, DUX4 protein^45^ and DUX4 downstream effects^28^.

Here we present a mathematical model of DUX4 expression in differentiated FSHD myoblasts, based on ordinary differential equations and stochastic gene expression. By analysing human myoblasts expressing inducible DUX4 as well as scRNAseq and snRNAseq of FSHD patient myocytes and myotubes, we compute experimental estimates for the parameters underlying our model. These include the first estimates of DUX4 transcription, translation and mRNA degradation rates. Simulating our model with experimentally derived parameters we find that it accurately predicts the proportion of DUX4 +ve/-ve and DUX4 target gene +ve/-ve cells observed in actual scRNAseq of FSHD patient myocytes. We package our model into graphical user interface tools to allow investigators to rapidly observe the impact of any given anti-DUX4 therapy on cell viability.

As with all models, ours are subject to assumptions, such as combining the multi-step processes of transcription and promoter activation into single steps. We also limited our model to differentiated muscle, when DUX4 expression is more robust. Thus, it is encouraging that we can simulate DUX4 and target gene expression single cell distributions which are indistinguishable from those observed in FSHD patients. Through theoretical investigation, we can also explore processes such as DUX4 protein syncytial diffusion and DUX4 target gene activation, which may be less accessible to experimentation. As more data on FSHD is generated our models will evolve, facilitating even more sophisticated understanding via estimation of further parameters.

DUX4 is myotoxic^27^, but how its rare expression drives and sustains significant pathology is unclear. DUX4 has been proposed to undergo a rare burst-like expression dynamic^24,26^, though no attempt has previously been made to understand DUX4 expression as a stochastic process. Here we model DUX4 and its target gene expression directly as stochastic processes via our promoter-switching model and estimate the underlying parameters. Our cell-biology informed model predicts that in single FSHD myocytes/myotubes DUX4 can drive significant cell death, eventually depleting the entire population, while being expressed in only 0.8% of live cells. This level matches well with the proportion of DUX4 positive cells seen in published studies^2,18,40^ and confirms that a burst-like mechanism of DUX4 expression can drive significant pathology while making DUX4 difficult to detect.

DUX4 has also been proposed to spread through the myofibre syncytium from its originator nucleus to ‘infect’ DUX4 naïve nuclei, bypassing the need for a rare expression burst and accelerating pathology^26,46^. The proportion of DUX4 target positive cells is greater in snRNAseq of syncytial FSHD myotubes than scRNAseq of unfused myocytes, supporting a syncytial diffusion mechanism. We model DUX4 in FSHD myotubes as an infectious agent, able to spread from one nucleus to another by adapting epidemiological compartment models, which we package as a cellular automaton to provide a realistic myotube surface on which to monitor DUX4 expression. Incorporation of a diffusion term is sufficient to account for the difference in DUX4 target gene +ve/-ve nuclear proportions, between syncytial FSHD myotubes and unfused FSHD myocytes. We thus demonstrate that the theoretical hypothesis of DUX4 syncytial diffusion is compatible with biological data.

Our model predicts <2.9% of single FSHD myocytes will be DUX4 target gene positive at any given time, despite significant DUX4 driven cell death. While in FSHD myotube nuclei this proportion can transiently spike at 70%, on average it remains much lower at only ^~^5.6%. This suggests that in FSHD muscle, expression of DUX4 target genes at any given time typically remains restricted to a small number of myonuclei. This may explain why DUX4 target gene expression is an inconsistent biomarker of FSHD muscle^20,21^. Moreover, our model predicts that DUX4 target gene expression does not typically increase over time, supporting the finding that DUX4 target gene expression in FSHD muscle has not validated as a marker of disease progression^47,48^. Lastly, due to the typically low levels observed, it may be difficult to detect changes in DUX4 target gene activation following anti-DUX4 therapies. This may have contributed to the recent phase 2b trial of losmapimod in FSHD failing to reach its primary outcome measure of suppressing DUX4 target gene expression in patient muscle biopsies^23^.

By modelling DUX4 target gene expression as stochastic processes, we found that DUX4 increases the proportion of time target gene promoters are in an active configuration, while curiously decreasing the normalised active promoter transcription rate. Through analytic investigation of our model, we found the rise in active promotor time drove increased DUX4 target gene expression, while the drop in normalised active promotor transcription rate dampened the noise associated with this higher expression level. This suggests that DUX4 modifies expression of its target genes to orchestrate a precise, stable activation. It has been shown that DUX4 is a pioneer factor and increases target gene promoter activation via C-terminal recruitment of p300 to drive H3K27 acetylation^29^. How DUX4 may limit normalised active promoter transcription rates of its target genes is unclear, but could involve interaction with the transcriptional complex, or feeding back to decrease the stability of target mRNA.

We thus present our model of DUX4 expression as a theoretical setting to understand the complex dynamics of this important disease gene and as an open source, in silico platform to rapidly and cheaply pre-screen anti-DUX4 therapy for FSHD.

## Methods

### Cell culture

DUX4 inducible immortalised human myoblasts LHCN-M2 were cultured in Promocell growth media (C-23060) supplemented with 15% FBS and 1:1000 Gentamycin (Sigma), in a humidified incubator at 5% CO_2_.

### RT-qPCR

Cells were grown in 6 well plates and total mRNA was isolated using the RNeasy Kit (Qiagen, 74-104) according to manufacturer’s instructions, after doxycycline stimulation. After reverse transcription with the Quantitect reverse transcription kit (Qiagen, 205311), SYBR green qPCR was performed (Takyon, UF-NSMT-B101) using a Viia7machine (ThermoFisher). DUX4 primers were: Fwd ACCTCTCCTAGAAACGGAGGC, Rev CAGCAGAGCCCGGTATTCTTC.

### Standard curve to compute d_0_

The length of the DUX4 construct used in our standard curve was 8938 bp and the number of copies per ng was calculated via the following formula:

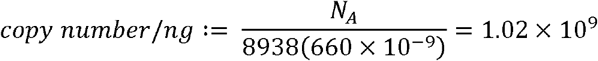

Where *N_A_* = 6.022 × 10^23^mol^-1^ is Avogadro’s number and 660 g/mol is the molecular weight of a single DNA base pair. Linear regression of the standard curve confirmed that DUX4 threshold cycles (Cts) are a linear function of log_10_(*copy number*) (R^2^=0.994), with slope 30.4 and intercept −2.70. These values were employed to convert DUX4 Cts from our iDUX4 assay to DUX4 copy number via the following formula:

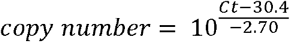

In our iDUX4 assay we sample RNA immediately after washing off doxycycline and at multiple time points post wash. We assume that no DUX4 can be transcribed post wash and model DUX4 copy number as an exponential decay:

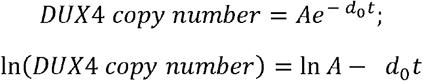

Linear regression of ln(*DUX4 copy number*) against time post wash (*t*) yielded a slope of 0.246, giving our estimate of *d*_0_, the degradation rate of DUX4.

### scRNAseq and snRNAseq data

Normalised read counts for scRNAseq of 7047 FSHD and control single myocytes produced by van den Heuvel et al.,^40^ were downloaded from the GEO data base accession GSE122873. This dataset contains 5133 single myocytes derived from 4 FSHD patients (2 FSHD1 and 2 FSHD2) and 1914 single myocytes from 2 control individuals. Myocytes were differentiated for 3 days in the presence of EGTA to prevent fusion.

Normalised read counts for snRNAseq of 317 FSHD and control single myonuclei produced by Jiang et al.,^16^ were downloaded from the GEO database accession GSE143492. This data comprised 47 FSHD2 nuclei at day 0 of differentiation 139 FSHD2 nuclei at day 3, from 2 sperate patients, alongside 54 control nuclei at day 0 and 77 control nuclei at day 3 from 2 separate individuals. For comparison with the scRNAseq data we focused on myonuclei from day 3 of differentiation.

All myocytes and myonuclei nuclei were assigned to one of 4 live cell compartments via the following criteria:

1. *S* – no detectable normalised reads for *DUX4, ZSCAN4, TRIM43, RFPL1, RFPL2, RFPL4B, PRAMEF1, PRAMEF2* and *PRAMEF12*.
2. *E* – at least one detectable normalised *DUX4* read, and no detectible normalised reads for *ZSCAN4, TRIM43, RFPL1, RFPL2, RFPL4B, PRAMEF1, PRAMEF2* and *PRAMEF12*.
3. *I* – at least one detectable normalised *DUX4* read, and at least one detectible normalised read for any of *ZSCAN4, TRIM43, RFPL1, RFPL2, RFPL4B, PRAMEF1, PRAMEF2* and *PRAMEF12*.
4. *R* – no detectable normalised reads for *DUX4*, and at least one detectible normalised read for any of *ZSCAN4, TRIM43, RFPL1, RFPL2, RFPL4B, PRAMEF1, PRAMEF2* and *PRAMEF12*.

All control myocytes/myonuclei were in the S compartment. The distribution of the 4 live cell compartments in the 5133 FSHD single myocytes and the 139 day 3 FSHD single myonuclei were compared via a Chi-squared test, significance was assessed at the 5% level.

### Estimation of MLEs of promoter-switching model parameters from scRNAseq data

To calculate transcription rate for DUX4 and DUX4 targets we adopt the approach employed by Larsson et al. and Trung et al. where a maximum likelihood estimation (MLE) methodology is used to infer the parameters for the two-state model of stochastic gene expression^32,49^. This is implemented in the *poisbeta* python package available at https://github.com/aksarkar/poisbeta. The procedure takes the Poisson-Beta distribution discussed in the results:

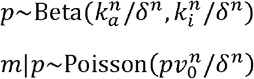

and minimizes the negative log-likelihood for each gene

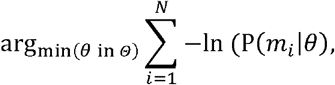

where *p* is a Beta-distributed variable describing the promotor state, *N* is the number of cells, *m_i_* is the number of mRNA transcripts for cell *i* = 1,…, *N*, and 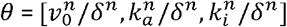, is the parameter space. From here on we will not include the normalising *δ^n^* term for brevity.

As a first estimate of *θ*, we use the method of moments derived by Peccoud and Ycart for the two-state promotor model^50^

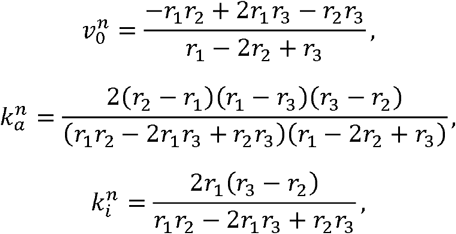

where

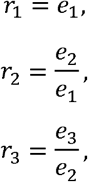

and

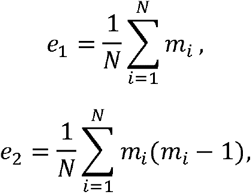

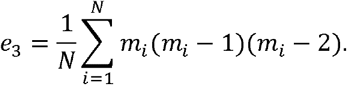

*θ* is then optimised via the Nelder-Mead method using Gauss-Jacobi quadrature to evaluate the likelihood function

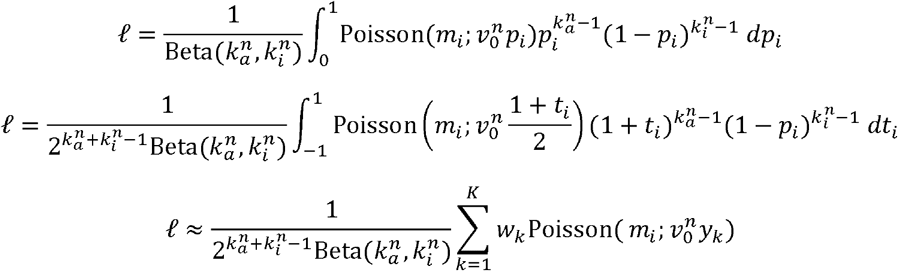

where *p_i_* = (*t_i_* + 1)/2 to substitute the beta-distributed parameter, *K* is the number of points integrated over, *w_k_* is the weight of the Jacobi polynomial of degree *k*, and *y_k_* is the root. The optimiser quadrature method, beta distribution, and Poisson distribution are implemented in the scipy package available from https://scipy.org/install/.

### Statistical comparison of promoter-switching model parameters

The three normalised parameters of the promoter-switching model 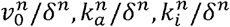, were computed as above for each of the 8 DUX4 target genes *ZSCAN4, TRIM43, RFPL1, RFPL2, RFPL4B, PRAMEF1, PRAMEF2* and *PRAMEF12*, in the 27/5133 FSHD single myocytes with detectable DUX4 mRNA and separately in the 5106 FSHD single myocytes without detectable DUX4, in the data set of van den Heuvel et al^40^.

The normalised active promoter transcription rate 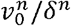, the proportion of time spent with promoter active 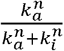 and the mean mRNA expression level *E*[*m*] for the 8 DUX4 target genes were compared between DUX4 positive and DUX4 negative myocytes via paired two-tailed Wilcoxon tests, significance was assessed at the 5% level.

### Hypothetical scenarios for raising E[m] under the promoter-switching model

We considered two hypothetical scenarios for raising *E*[*m*] under the promotor-switching model to observe the impact on *Var*[*m*]. In the first 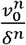 is increased while keeping other parameters constant in the second 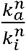 is increased keeping other parameters constant. Simple algebraic manipulation allows analytical solutions for the raised parameters and thus *Var*[*m*].

In the case of 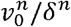, we assume that 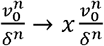 for some *x* > 1, the rise in *E*[*m*] in this case is simply:

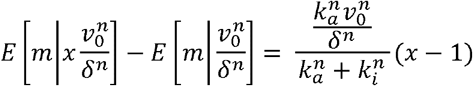

We can thus choose *x*, to achieve the rise in *E*[*m*] seen under DUX4 expression *d*(*μ*) as:

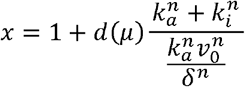

Which can be substituted into the formula for *Var*[*m*] to compute 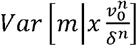 giving the expected variance of the mRNA copy distribution if *E*[*m*] is increased by *d*(*μ*), solely by increasing 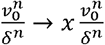 for some *x* > 1.

In the case of 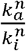, we assume that 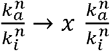, we further assume that the rise in active transition rate is mirrored by a drop in the inactivation rate, i.e., the two parameters are not mutually exclusive, thus: 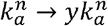 and 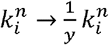, where *y*^2^ = *x*. The rise in *E*[*m*] in this case is:

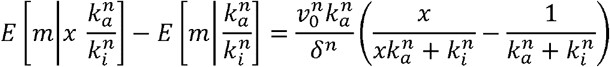

We can thus choose *x*, to achieve the rise in *E*[*m*] seen under DUX4 expression *d*(*μ*) as:

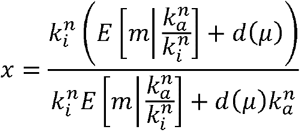

For each of the 8 DUX4 target genes we considered the parameters of the promoter switching model 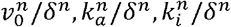, computed over the 5106 DUX4 -ve FSHD single myocytes from the data set of van den Heuvel et al., as the starting parameters to input into the above formulae for *x*. Comparing the mean expression of each target in the 27 DUX4 +ve myocytes and the 5106 DUX4 -ve myocytes we computed *d*(*μ*). For each target we thus computed 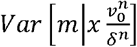 and 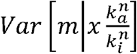, the target mRNA copy number variances under our 2 hypothetical scenarios. These variances were compared with the true mRNA copy number variance in the 27 DUX4 +ve myocytes, as well as with each other via two tailed paired Wilcoxon tests, significance was assessed at the 5% level.

### Compartment model simulation

After estimation of the 5 parameters *V_D_, d*_0_, *V_T_, T_D_* and *D_r_*, the ODEs underlying the compartment model **(Figure 1B)** were simulated from an initial state of all cells in state *S*(0), over a period of 100 days in timesteps of 1 hour using the deSolve package in R^51^. For comparison with the 5133 single FSHD myocytes differentiated for 3 days described by van den Heuvel et al.,^40^, an initial population sized 5133 was simulated for 3 days, after which 6.901% of starting cells were in the dead cell state. In order to replicate the live cell population of 5133 after 3 days we thus employed a starting population of 5133(1+0.06901) cells in state *S*(0). After simulation for 3 days the proportion of cells in each live cell state *S*(3 *days*), *E*(3 *days*), *I*(3 *days*) and *R*(3 *days*) was compared between the model simulation and the true scRNAseq data, via a Chi-squared test.

For the syncytial compartment model, simulation was performed as above with *V_D_, d*_0_, *V_T_, T_D_* and *D_r_* unchanged. The additional parameter Δ was estimated via genetic algorithm (below) alongside the starting population *n* of cells in state *S*. After simulation for 2 days (as there is assumed no syncytium for the first 24 hours of fusion) the proportion of cells in each live cell state was compared between the model simulation and the true snRNAseq data of Jiang et al.,^16^ via a Chi-Squared test. Significance was assessed at the 5% level.

### Cellular Automaton Model

The cellular automaton model was evolved on an *l* × *c* rectangular grid describing the length and circumference of myotube. Individual cells on the grid correspond to single myonuclei in a syncytium. Each cell was evolved stochastically over a discrete time-step of one hour. In each timestep 2 independent actions were performed.

First the state of each cell was evolved according to the non-syncytial compartment model. Transition from connected states was modelled as an exponential process with rate parameter equal to the transition rate (e.g., transition from *S* to *E* occurred according to exp (*V_D_*)). An event occurring or not under this exponential distribution within 1 hour was simulated during each time step. If the event occurred a transition in cell state occurred, otherwise the cell state remained the same. In the situation where a cell state could transition in 2 possible directions (e.g., state *E* can transition to *S* at rate *d*_0_ and to *I* at rate *T_D_V_T_*) competing exponential clocks were set up. Using the property that min(exp(*A*), exp(*B*))~exp (*A* + *B*), we simulated a transition occurring in either direction within 1 hour. If a transition occurred, we simulated the exponential clocks describing transition in either direction independently, and transitioned the cell state according to whichever experienced an event faster. If no transition occurred the cell state remained the same.

Once every cell was evolved according the non-syncytial compartment model we updated the model according to diffusion of DUX4. Cells in states *I* or *R* at the start of the time-step can interact with cells in states *S* or *E* if they are among their 8 immediate neighbours **(Figure 6B).** Each interaction was again modelled as an exponential distribution with rate Δ, which was simulated for an event occurring within 1 hour. If the event occurred then the state of the neighbouring cell was updated *S* → *I* or *E* → *R*, otherwise the state of the neighbouring cell remained the same.

We implement a boundary condition so that that cells at position 1 on the circumference dimension are able to interact with cells in position *c*, to allow our grid topological equivalence to a cylinder **(Figure 6A).**

The cellular automaton model was evolved from a starting distribution of all cells in state S over 48 hour time-steps, employing the parameters *V_D_, d*_0_, *V_T_, T_D_* and *D_r_* estimated from the non-syncytial compartment model and Δ, *l* and *c* estimated from genetic algorithm and clustering analysis (see below). Due to the stochastic nature of the exponential clocks involved, 100 simulations were performed to obtain an average behaviour of cell type proportions under our model. The proportion of cells in each of the live cell states obtained from the average behaviour of our model was compared to the true snRNAseq data of Jiang et al.,^16^ via a Chi-Squared test. Significance was assessed at the 5% level.

### Genetic Algorithms

We employed genetic algorithms twice, firstly in the syncytial compartment model to fit the number of myonuclei *n* in the starting state *S* and the diffusion parameter Δ. Secondly in the cellular automaton model to again fit Δ, and the length *l* and circumference *c* of the automaton grid, and thus again the number of cells *n* = *l* × *c* in the starting state *S*.

Both algorithms employed the same fitness function aimed at minimising the difference in the proportion of cells in the live states between the simulated model after 2 days and the snRNAseq of Jiang et al^16^:

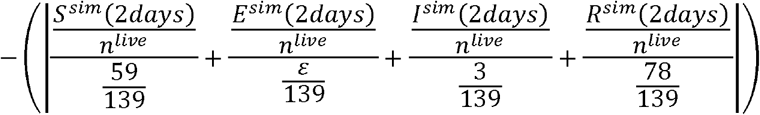

Where *n^live^* ≔ *n* – *D^sim^*(2*days*) and *ε* ≪ 1 is chosen to prevent singularity while rewarding models which obtain values of 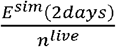 close to 0. This fitness function was chosen as it outperformed conventional functions based on Minkowski and Euclidean distances. A value of *ε* = 0.1, proved sufficient to generate models with live cell state proportions indistinguishable from the snRNAseq data via Chi-squared test.

Genetic algorithms were implemented using the GA package in R^52^ using default parameters: starting population of 50 models, elitism of 5%, mutation probability 0.1, crossover probability 0.8, an optimal model was selected if maximum fitness was stable for 10 generations.

The cellular automaton model evolves stochastically and thus an optimal fitness function for a population could represent an optimal parameter regime, or an unlikely simulation of a sub-optimal parameter regime. We thus ran 100 genetic algorithms on the cellular automaton to generate 100 optimal parameter regimes and examined their structure via clustering analysis (below).

### Clustering analysis

Clustering analysis was performed on the 3×100 feature space of parameter regimes Δ, *l* and *c* found optimal in the 100 genetic algorithms performed on the cellular automaton model. The ConsensusClusterPlus package in R^53^ implemented K-medoids clustering using a Euclidean distance metric, and consensus-CDF cluster stability plots ascertained the optimal number of clusters in the parameter feature space as 5.

Two-tailed paired Wilcoxon tests comparing the distribution of *l* and *c* values in each cluster, demonstrated that only one parameter cluster output significantly long and thin myotubes, in line with the microanatomy of muscle **(Figure 6D).** The average values of Δ, *l* and *c* from this cluster were thus considered the optimal parameter regime for the cellular automaton model.

### Shiny tools

Three GUI tools were written using the shiny package in R^54^. The first tool outputs the compartment models for user parameter inputs and is implemented using the deSolve package^51^. The second tool outputs a dynamic realisation of the stochastic cellular automaton model for user parameter inputs, the automaton is displayed using the lattice package in R^55^. The third tool displays the proportion of dead cells under the compartment model comparing our experimentally derived parameter regime to a user selected parameter regime. In addition the third tool simulates 2 realisations of the cellular automaton model, one using our experimentally derived parameter regime and another using the user selected regime (*l* and *c* are kept the same for both simulations to allow comparable starting populations). Survival analysis using Cox-proportional hazard models are implemented via the survival package in R^56^, to compare cell death rates in the two realisations, with *p*-values displayed on a corresponding Kaplan-Meier plot.

The 3 tools can be accessed at:

4. Compartment Models: https://crsbanerji.shinyapps.io/compartment_models/
5. Cellular Automaton: https://crsbanerji.shinyapps.io/ca_shiny/
6. Survival Analysis: https://crsbanerji.shinyapps.io/survival_sim/

## Acknowledgements

MVC was supported by the EPSRC Centre for Doctoral Training in Sustainable Chemical Technologies (EP/L016354/1) and Friends of FSH research (Project: An in-silico approach to understanding DUX4 expression). JP was supported by Muscular Dystrophy UK (19GRO-PG12-0493) and currently by the FSHD Society (FSHD-Winter2021-4491649104). MG is supported by the Medical Research Council (MR/S002472/1). The Zammit lab was also generously supported by Association Française contre les Myopathies.

## References

1. Deenen, J. C. W. et al. Population-based incidence and prevalence of facioscapulohumeral dystrophy. Neurology 83, 1056–1059 (2014).

2. Banerji, C. R. S. & Zammit, P. S. Pathomechanisms and biomarkers in facioscapulohumeral muscular dystrophy: roles of DUX4 and PAX7. EMBO Mol. Med. 13, e13695 (2021).

3. Dahlqvist, J. R. et al. Evaluation of inflammatory lesions over 2 years in facioscapulohumeral muscular dystrophy. Neurology 10.1212/WNL.0000000000010155 (2020). doi: 10.1212/WNL.0000000000010155

4. Tawil, R., Storvick, D., Feasby, T. E., Weiffenbach, B. & Griggs, R. C. Extreme variability of expression in monozygotic twins with FSH muscular dystrophy. Neurology 43, 345–348 (1993).

5. Banerji, C. R. S. et al. Facioscapulohumeral muscular dystrophy 1 patients participating in the UK FSHD registry can be subdivided into 4 patterns of self-reported symptoms. Neuromuscul. Disord. 315–328 (2020). doi:10.1016/j.nmd.2020.03.001

6. Schepelmann, K. et al. Socioeconomic burden of amyotrophic lateral sclerosis, myasthenia gravis and facioscapulohumeral muscular dystrophy. J. Neurol. 257, 15–23 (2010).

7. Lemmers, R. J. L. F. et al. A unifying genetic model for facioscapulohumeral muscular dystrophy. Science 329, 1650–1653 (2010).

8. Lemmers, R. J. L. F. et al. Digenic inheritance of an SMCHD1 mutation and an FSHD-permissive D4Z4 allele causes facioscapulohumeral muscular dystrophy type 2. Nat. Genet. 44, 1370–1374 (2012).

9. van den Boogaard, M. L. et al. Mutations in DNMT3B Modify Epigenetic Repression of the D4Z4 Repeat and the Penetrance of Facioscapulohumeral Dystrophy. Am. J. Hum. Genet. 98, 1020–1029 (2016).

10. Hamanaka, K. et al. Homozygous nonsense variant in LRIF1 associated with facioscapulohumeral muscular dystrophy. Neurology 94, e2441–e2447 (2020).

11. De Iaco, A. et al. DUX-family transcription factors regulate zygotic genome activation in placental mammals. Nat. Genet. 49, 941–945 (2017).

12. Tawil, A. et al. Design of a Phase 2, Randomized, Double-Blind, Placebo-Controlled, 24-Week, Parallel-Group Study of the Efficacy and Safety of Losmapimod in Treating Subjects with Facioscapulohumeral Muscular Dystrophy (FSHD): ReDUX4 (1592). Neurology 94, (2020).

13. Gall, L. Le, Sidlauskaite, E., Mariot, V. & Dumonceaux, J. Therapeutic Strategies Targeting DUX4 in FSHD. J. Clin. Med. 9, 1–20 (2020).

14. Jones, T. I. et al. Facioscapulohumeral muscular dystrophy family studies of DUX4 expression: evidence for disease modifiers and a quantitative model of pathogenesis. Hum. Mol. Genet. 21, 4419–4430 (2012).

15. Beermann, M. Lou, Homma, S. & Miller, J. B. Proximity ligation assay to detect DUX4 protein in FSHD1 muscle: a pilot study. BMC Res. Notes 15, 1–6 (2022).

16. Jiang, S. et al. Single-nucleus RNA-seq identifies divergent populations of FSHD2 myotube nuclei. PLOS Genet. 16, e1008754 (2020).

17. Van Den Heuvel, A. et al. Single-cell RNA sequencing in facioscapulohumeral muscular dystrophy disease etiology and development. Hum. Mol. Genet. 28, 1064–1075 (2019).

18. Snider, L. et al. Facioscapulohumeral Dystrophy: Incomplete Suppression of a Retrotransposed Gene. PLoS Genet. 6, e1001181 (2010).

19. Rickard, A. M., Petek, L. M. & Miller, D. G. Endogenous DUX4 expression in FSHD myotubes is sufficient to cause cell death and disrupts RNA splicing and cell migration pathways. Hum. Mol. Genet. 24, 5901–5914 (2015).

20. Banerji, C. R. S. et al. PAX7 target genes are globally repressed in facioscapulohumeral muscular dystrophy skeletal muscle. Nat. Commun. 8, 2152 (2017).

21. Banerji, C. R. S. & Zammit, P. S. PAX7 target gene repression is a superior FSHD biomarker than DUX4 target gene activation, associating with pathological severity and identifying FSHD at the single-cell level. Hum. Mol. Genet. (2019).

22. Banerji, C. R. S., Panamarova, M. & Zammit, P. S. DUX4 expressing immortalized FSHD lymphoblastoid cells express genes elevated in FSHD muscle biopsies, correlating with the early stages of inflammation. Hum. Mol. Genet. 29, 2285–2299 (2020).

23. Jagannathan, S. et al. Meeting report: the 2021 FSHD International Research Congress. Skelet. Muscle 12, 1–10 (2022).

24. Bosnakovski, D. et al. Muscle pathology from stochastic low level DUX4 expression in an FSHD mouse model. Nat. Commun. 8, 1–9 (2017).

25. Bosnakovski, D. et al. Transcriptional and cytopathological hallmarks of FSHD in chronic DUX4-expressing mice. J. Clin. Invest. 130, 2465–2477 (2020).

26. Tassin, A. et al. DUX4 expression in FSHD muscle cells: How could such a rare protein cause a myopathy? J. Cell. Mol. Med. 17, 76–89 (2013).

27. Kowaljow, V. et al. The DUX4 gene at the FSHD1A locus encodes a pro-apoptotic protein. Neuromuscul. Disord. 17, 611–623 (2007).

28. Heher, P. et al. Interplay between mitochondrial reactive oxygen species, oxidative stress and hypoxic adaptation in facioscapulohumeral muscular dystrophy: Metabolic stress as potential therapeutic target. Redox Biol. 51, 102251 (2022).

29. Choi, S. H. et al. DUX4 recruits p300/CBP through its C-terminus and induces global H3K27 acetylation changes. Nucleic Acids Res. 44, 5161–5173 (2016).

30. Knopp, P. et al. DUX4 induces a transcriptome more characteristic of a less-differentiated cell state and inhibits myogenesis. J. Cell Sci. 129, 3816–3831 (2016).

31. Shahrezaei, V. & Swain, P. S. Analytical distributions for stochastic gene expression. Proc. Natl. Acad. Sci. U. S. A. 105, 17256–17261 (2008).

32. Vu, T. N. et al. Beta-Poisson model for single-cell RNA-seq data analyses. Bioinformatics 32, 2128–2135 (2016).

33. Kim, J. K. & Marioni, J. C. Inferring the kinetics of stochastic gene expression from single-cell RNA-sequencing data. Genome Biol. 14, 1–12 (2013).

34. Ganassi, M., Figeac, N., Reynaud, M., Ortuste Quiroga, H. P. & Zammit, P. S. Antagonism Between DUX4 and DUX4c Highlights a Pathomechanism Operating Through β-Catenin in Facioscapulohumeral Muscular Dystrophy. Front. Cell Dev. Biol. 10, 1419 (2022).

35. Yang, E. et al. Decay Rates of Human mRNAs: Correlation With Functional Characteristics and Sequence Attributes. Genome Res. 13, 1863–1872 (2003).

36. Young, J. M. et al. DUX4 binding to retroelements creates promoters that are active in FSHD muscle and testis. PLoS Genet. 9, e1003947 (2013).

37. Young, J. M. et al. DUX4 Binding to Retroelements Creates Promoters That Are Active in FSHD Muscle and Testis. PLoS Genet. 9, (2013).

38. Geng, L. N. et al. DUX4 activates germline genes, retroelements, and immune mediators: implications for facioscapulohumeral dystrophy. Dev. Cell 22, 38–51 (2012).

39. Banerji, C. R. S. et al. Dynamic transcriptomic analysis reveals suppression of PGC1α/ERRα drives perturbed myogenesis in facioscapulohumeral muscular dystrophy. Hum. Mol. Genet. (2018). doi:10.1093/hmg/ddy405

40. van den Heuvel, A. et al. Single-cell RNA-sequencing in facioscapulohumeral muscular dystrophy disease etiology and development. Hum. Mol. Genet. (2018). doi:10.1093/hmg/ddy400

41. Gall, L. Le, Sidlauskaite, E., Mariot, V. & Dumonceaux, J. Therapeutic Strategies Targeting DUX4 in FSHD. J. Clin. Med. 2020, Vol. 9, Page 2886 9, 2886 (2020).

42. Block, G. J. et al. Wnt/β-catenin signaling suppresses DUX4 expression and prevents apoptosis of FSHD muscle cells. Hum. Mol. Genet. 22, 4661–4672 (2013).

43. Wallace, L. M. et al. RNA Interference Inhibits DUX4-induced Muscle Toxicity In Vivo: Implications for a Targeted FSHD Therapy. Mol. Ther. 20, 1417–1423 (2012).

44. Ciszewski, L. et al. G-quadruplex ligands mediate downregulation of DUX4 expression. Nucleic Acids Res. 48, 4179–4194 (2020).

45. Klingler, C. et al. DNA aptamers against the DUX4 protein reveal novel therapeutic implications for FSHD. FASEB J. 34, 4573–4590 (2020).

46. Barro, M. et al. Myoblasts from affected and non-affected FSHD muscles exhibit morphological differentiation defects. J. Cell. Mol. Med. 14, 275–289 (2010).

47. Banerji, C. R. S. PAX7 target gene repression associates with FSHD progression and pathology over 1 year. Hum. Mol. Genet. 00, (2020).

48. Wong, C.-J. et al. Longitudinal measures of RNA expression and disease activity in FSHD muscle biopsies. Hum. Mol. Genet. 00, (2020).

49. Larsson, A. J. M. et al. Genomic encoding of transcriptional burst kinetics. Nature 565, 251–254 (2019).

50. Peccoud, J. & Ycart, B. Markovian Modeling of Gene-Product Synthesis. Theor. Popul. Biol. 48, 222–234 (1995).

51. Soetaert, K., Petzoldt, T. & Setzer, R. W. Solving Differential Equations in R: Package deSolve. J. Stat. Softw. 33, 1–25 (2010).

52. Scrucca, L. GA: A Package for Genetic Algorithms in R. J. Stat. Softw. 53, 1–37 (2013).

53. Wilkerson, M. D. & Hayes, D. N. ConsensusClusterPlus: a class discovery tool with confidence assessments and item tracking. Bioinformatics 26, 1572–3 (2010).

54. Chang, W., Cheng, J., Allaire, J., Xie, Y. & McPherson, J. shiny: Web Application Framework for R. R package version 1.2.0. Https://CRAN.R-project.org/package=shiny, (2014).

55. Sarkar & Deepayan. Lattice: Multivariate data visualization with R. R package version 0.20-38. (2008).

56. Therneau, T. M. & Grambsch, P. M.Modeling survival data⍰: extending the Cox model. (2018).

